# Genetic diversity of *bla*_KPC_-gene-containing IncF plasmids from epidemiologically related and unrelated Enterobacteriaceae

**DOI:** 10.1101/2020.11.01.362400

**Authors:** Joep J.J.M. Stohr, Marjolein F. Q. Kluytmans-van den Bergh, Veronica A.T.C. Weterings, John W. A. Rossen, Jan A. J. W. Kluytmans

## Abstract

**Background:** Limited information is available on whether *bla*_KPC_-containing plasmids from isolates in a hospital outbreak can be differentiated from epidemiologically unrelated *bla*_KPC_-containing plasmids based on sequence data.

**Objective:** This study aimed to evaluate the performance of three approaches to distinguish epidemiologically related from unrelated *bla*_KPC_-containing IncF plasmids.

**Method:** Epidemiologically related isolates, were short- and long-read whole genome sequenced on an Illumina MiSeq and MinION sequencer. A hybrid assembly was performed and plasmid sequences were extracted from the assembly graph. Epidemiologically unrelated plasmid sequences were extracted from the GenBank. Pairwise comparisons were performed of epidemiologically related and unrelated plasmids based on SNP differences using snippy, phylogenetic distance using Roary and using a similarity index that penalizes size differences between plasmids (Stoesser-index). The percentage of pairwise comparisons misclassified as genetically related or as clonally unrelated was determined using different genetic thresholds for genetic relatedness for all three comparison methods.

**Results:** Despite the median number of SNP differences, Roary phylogenetic distance, and Stoesser-index differed between the epidemiologically related and unrelated plasmids, the range of differences overlapped between the two comparison groups for all three comparison methods. When using a genetic similarity threshold that classified 100% of epidemiologically related plasmid pairs as genetically related, the percentages of plasmids misclassified as epidemiologically related ranged from 6.7% (Roary) to 20.8% (Stoesser-index).

**Discussion:** Although epidemiologically related plasmids can be distinguished from unrelated plasmids based on genetic similarity, epidemiologically related and unrelated *bla*_KPC_-containing IncF plasmids show a high degree of sequence similarity. The phylogenetic distance as determined using Roary showed the highest degree of discriminatory power between the epidemiologically related and unrelated plasmids.

**Impact statement:** Accurately distinguishing epidemiologically related from unrelated plasmids is essential to detect nosocomial plasmid transmission in outbreaks. However, limited information is available on whether *bla*_KPC_-containing plasmids from isolates in a hospital outbreak can be differentiated from epidemiologically unrelated *bla*_KPC_-containing plasmids based on sequence data. This study aimed to evaluate the performance of three approaches to distinguish epidemiologically related from unrelated *bla*_KPC_-containing IncF plasmids. Pairwise comparisons were performed of epidemiologically related and unrelated plasmids based on SNP differences using snippy, phylogenetic distance using Roary and using a similarity index that penalizes size differences between plasmids (Stoesser-index). Based on our results, epidemiologically related plasmids can be distinguished from unrelated plasmids based on genetic similarity. Despite this, epidemiologically related and unrelated *bla*_KPC_-containing IncF plasmids show a high degree of sequence similarity and judgements on the horizontal transfer of these plasmids during hospital outbreaks based on genetic identity should be made with caution. The phylogenetic distance determined using Roary showed the highest discriminatory power between the epidemiologically related and unrelated plasmids.

**Data summary:** Short-and long-read sequence data of the epidemiologically related Enterobacteriaceae isolates included in this study are available from the publicly available European Nucleotide Archive of the European Bioinformatics Institute under study accession number: PRJEB41009. The authors confirm that all supporting data have been provided within the article and through the supplementary data files.

## Introduction

Infections with carbapenem-resistant Enterobacteriaceae are associated with increased mortality and have emerged as an urgent public health threat (^1–3^). Worldwide, one of the most frequent mechanisms for carbapenem resistance in Enterobacteriaceae is the production of *K. pneumoniae* carbapenemase (KPC) (^4^). Nosocomial transmission plays an important role in spreading these KPC-producing Enterobacteriaceae, and several hospital outbreaks have been described (^5–8^). Molecular typing is often used to detect the source and route of transmission in these hospital outbreak settings (^9^). Whole-genome sequencing of bacterial isolates is currently the ultimate tool for molecular typing, enabling comparison of the entire bacterial chromosome to identify bacterial clones with great precision using either a gene-by-gene or single nucleotide polymorphism (SNP) approach (^10^). However, the gene encoding KPC-production, *bla*_KPC_, is not located on the bacterial chromosome but on large conjugative resistance plasmids. These plasmids can transfer between isolates, and nosocomial transmission of these plasmids can go undetected when only molecular typing of the bacterial chromosome is performed (^11^). Several reports have already described plasmid spread between different isolates and patients during hospital outbreaks (^12–16^). Analysing these resistance plasmids is typically performed by generating a combination of short- and long-read sequence data, enabling hybrid assembly algorithms to perform complete plasmid assemblies (^14,17–19^). Accurately distinguishing epidemiologically related from unrelated plasmids (e.g., based on the number of SNP differences) is essential to detect or exclude nosocomial plasmid transmission in outbreaks. However, limited information is available on whether *bla*_KPC_-containing plasmids from isolates in a hospital outbreak can be differentiated from epidemiologically unrelated *bla*_KPC_-containing plasmids based on sequence data. This study aimed to evaluate the performance of three approaches based on determining SNP differences, phylogenetic distance, and a similarity index, to distinguish epidemiologically related from unrelated *bla*_KPC_-containing IncF plasmids.

## Methods

### Outbreak and plasmid selection

From June to December 2013, an outbreak occurred in an 800-bed teaching hospital (approximately 40,000 admissions/year) and a 150-bed nursing home in Breda, the Netherlands. In total, six patients were colonised or infected with KPC-producing Enterobacteriaceae belonging to three different species (*K. pneumoniae, K. aerogenes*, and *E. coli*). The outbreak was comprehensively described by Weterings et al. (^6^). Epidemiologically related isolates (i.e., belonging to the outbreak) were selected in such a way that at least every unique *bla*_KPC_ gene containing isolate, based on species identification, antimicrobial susceptibility testing, and molecular typing (^6^), was included. Moreover, one additional KPC-producing *K. pneumoniae* isolate was included that was cultured after the outbreak period in February 2014 in a rectal swab taken from patient 3. The antimicrobial susceptibility testing of this isolate differed from the outbreak KPC-producing *K. pneumoniae* isolates, being measured susceptible to ciprofloxacin, tobramycin, and trimethoprim/sulfamethoxazole using Vitek-2 (bioMérieux, Marcy-l’Étoile, Franc) automated susceptibility testing and using EUCAST breakpoints v10.0. Epidemiologically unrelated plasmid sequences were extracted from the GenBank on 09-11-2017. All plasmid sequences containing a *bla*_KPC*-2*_ gene that contained the replicons IncFII_K2_-FIB but isolated in different countries or a different year were extracted and included in the study.

### Whole-genome sequencing and analysis

Epidemiologically related isolates were short-read sequenced on an Illumina MiSeq using Nextera XT chemistry generating 250-bp paired-end reads (Illumina, San Diego, United States) and long-read sequenced on a MinION sequencer using the FLO-MIN106D flow cell and the Rapid Barcoding Sequencing Kit SQK RBK004 according to the standard protocol provided by the manufacturer (Oxford Nanopore Technologies, Oxford, United Kingdom). A hybrid assembly of long-read and short-read sequence data was performed using Unicycler v.0.8.4 (^21^). Whole-genome MLST (wgMLST) (core and accessory genome) was performed for all sequenced isolates using Ridom SeqSphere+, version 4.1.9 (Ridom, Münster, Germany). Species-specific typing schemes were used as described by Kluytmans-van den Bergh et al. (^22^). The pairwise genetic difference between isolates of the same species was calculated by dividing the number of allele differences by the total number of alleles shared between the two sequences in the wgMLST typing scheme. The sequenced isolates’ genomes were uploaded to the online bioinformatic tools ResFinder v.3.1 and PlasmidFinder v.2.0 (Center for Genomic Epidemiology, Technical University of Denmark, Lyngby, Denmark) (^23,24^). Acquired resistance genes were called when at least 60% of the length of the best matching gene in the ResFinder database was covered with a sequence identity of at least 90%. Plasmid replicon genes were called when at least 60% of the sequence length of the replicon gene in the PlasmidFinder database was covered with a sequence identity of at least 80%.

### Plasmid analysis

Circular components created by the hybrid assembly that were smaller than 1000kb and contained a *bla*_KPC_-gene were extracted from the assembly graph using BANDAGE v0.8.1 (^25^). All extracted plasmid components and the plasmid sequences extracted from the GenBank were annotated using Prokka v1.13.3 (^26^). The number of SNP differences between the plasmids were calculated using snippy v4.4.5 with plasmid pKpQIL (GenBank accession number: NC_014016.1) as the reference sequence. A pangenome was constructed, and phylogenetic distance based on gene-presence/-absence between the different plasmids was determined using Roary v1.13.2 (^27^). All plasmids were pairwise aligned using *dnadiff*, and a similarity index was calculated between the different plasmids as described by Stoesser et al. (^20^) (Stoesser-similarity index). This similarity index can vary from 0 (completely unsimilar plasmids) to 1 (identical plasmids) and, contrary to both Roary phylogenetic distance and number of SNP differences, penalises size differences between plasmids. Pairwise comparisons of SNP differences, Roary phylogenetic distance, Stoesser-similarity were performed between: 1) epidemiologically related plasmids and 2) the first plasmid isolated from the index patient in the outbreak and the epidemiologically unrelated plasmid sequences extracted from GenBank. The percentage of pairwise comparisons misclassified as genetically related or as clonally unrelated was determined using different genetic thresholds for genetic relatedness for all three comparison methods: SNP thresholds tested ranged from 0 to 50 with steps of 1, Roary thresholds from 0.0 to 0.5 with steps of 0.01 and Stoesser index thresholds from 0.5 to 1 with steps of 0.00001.

### Statistical analysis

The median number of SNP differences, the median Roary phylogenetic distance, and the median Stoesser-similarity index were compared between epidemiologically related and unrelated plasmids using Mann-Whitney U tests (SciPy v1.5.0). Spearman’s rank correlation coefficients between the number of SNP differences, Roary phylogenetic distance and the Stoesser-similarity index were calculated using SciPy v1.5.0.

### Accession numbers

Generated raw reads were submitted to the European Nucleotide Archive (ENA) of the European Bioinformatics Institute (EBI) under the study accession number: PRJEB41009.

## Results

The percentage of wgMLST allele differences between the 3 *K. pneumoniae* isolates ranged from 0.002% (Pk1 vs. Pk2) to 0.770% (Pk1 (and Pk2) vs. Pk4). The percentage of wgMLST allele differences between the 2 *K. aerogenes* isolates was 0.979%. In all epidemiologically-related isolates, a circular plasmid contig containing a *bla*_KPC*-2*_ gene (located within a Tn*4401a* transposon) and an IncFII(k2) and IncFIB plasmid replicon was detected and extracted from the assembly graph (**Table 1**). The plasmid contigs were either 113638 or 113639 base pairs in size, had a GC-content of 53.9%, and contained the acquired beta-lactam resistance genes *bla*_TEM1A_ and *bla*_OXA-9_. Fifteen epidemiologically unrelated plasmids were detected in and extracted from GenBank. The plasmids were isolated from patients in 5 different countries (Greece, Italy, USA, Australia, and the UK) between 2007 and 2017 and ranged in size from 99142 to 117916 base pairs (**Table 1**) (^28–30^).

**Table 1.** Plasmids included from the outbreak and GenBank.

| Patient | Host species | Host MLST | Name plasmid | Epidemiologically related | Accession number | Year isolated | Location | Plasmid size | n of contigs plasmid |
| --- | --- | --- | --- | --- | --- | --- | --- | --- | --- |
| P1 (index) | klpne | 258 | Pk1 | yes | n.a. | 2013 | Netherlands, Breda | 113639 | 1 |
| P2 | klpne | 258 | Pk2 | yes | n.a. | 2013 | Netherlands, Breda | 113638 | 1 |
| P3 | klaer | n.a. | Pk3 | yes | n.a. | 2013 | Netherlands, Breda | 113639 | 1 |
| P3 | escol | 2598 | Pe1 | yes | n.a. | 2013 | Netherlands, Breda | 113639 | 1 |
| P3 | klpne | 309 | Pk4 | yes | n.a. | 2014 | Netherlands, Breda | 113639 | 1 |
| P5 | klaer | n.a. | Pk5 | yes | n.a. | 2013 | Netherlands, Breda | 113639 | 1 |
| n.a. | klpne | 258 | pKP1504-kpc | no | NC_023903.1 | 2008 | Greece, Athens | 113640 | n.a. |
| n.a. | klpne | 147 | pKP1780-kpc | no | NC_023904.1 | 2009 | Greece, Heraklion | 113622 | n.a. |
| n.a. | klpne | 234 | pKpQil-234 | no | NC_025187.1 | 2009 | USA, Central New Jersey | 114464 | n.a. |
| n.a. | klpne | 10 | pKpQil-10 | no | NC_025166.1 | 2010 | USA, New York City (New York) | 113639 | n.a. |
| n.a. | klpne | 35 | pKP3913-kpc | no | NC_023906.1 | 2011 | Greece, Athens | 113640 | n.a. |
| n.a. | klpne | 11 | pKP1870-kpc | no | NC_023905.1 | 2009 | Greece, Agios Niokolaos | 116047 | n.a. |
| n.a. | escol | 131 | pKpQIL-Ec | no | NC_025167.1 | 2010 | USA, Northen New Jersey | 99142 | n.a. |
| n.a. | klpne | 258 | pKpQIL-531 | no | NZ_CP008833.1 | 2013 | USA, Bethesda (Maryland) | 113639 | n.a. |
| n.a. | klpne | 258 | pUHKPC07 | no | NZ_CP011986.1 | 2007 | USA, Cleveland (Ohio) | 113639 | n.a. |
| n.a. | klpne | 258 | pUHKPC33 | no | NZ_CP011991.1 | 2008 | USA, Cleveland (Ohio) | 113638 | n.a. |
| n.a. | klpne | 258 | p500_1420 | no | NZ_CP011981.1 | 2012 | USA, North Eastern Ohio | 130552 | n.a. |
| n.a. | klpne | 258 | CR14_p3 | no | NZ_CP015395.1 | 2012 | USA, Valhalla (New York) | 116419 | n.a. |
| n.a. | klpne | n.a. | KPN207_p2 | no | NZ_LT216438.1 | 2016* | UK | 117916 | n.a. |
| n.a. | klpne | 258 | pAUSMDU8079-2 | no | NZ_CP022693.1 | 2017* | Australia | 113639 | n.a. |
| n.a. | Klpne | 258 | pIT-01C03 | no | HG969995 | 2011 | Italy, Milan | 113642 | n.a. |

\*Year/location of GenBank submission, no further information available; klpne: *K. pneumoniae*; kleaer: *K. aerogenes*; escol : *E. coli*.

The number of pairwise comparisons was 15 for the epidemiologically related plasmids and 120 for the epidemiologically unrelated plasmids (including the plasmid from the index patient) (**Table 2**). The median number of SNP differences varied significantly between the two groups (p < 0.001) (**Table 2**). However, the range of SNP differences overlapped, ranging from 0 to 1 for the epidemiologically related plasmids and from 0 to 674 for the epidemiologically unrelated plasmids (**Table 2; Supplementary Table S1**). A total number of 192 genes were detected, of which 56 were present in all plasmids. The phylogenetic distance as determined using Roary ranged from 0.00 to 0.06 between the epidemiologically related plasmids and from 0.00 to 1.69 between the epidemiologically unrelated plasmids (p < 0.001) (**Table 2; Supplementary Table S2**). The Stoesser-similarity index ranged from 0.99 to 1 between the epidemiologically related plasmids and from 0.51 to 1 between the epidemiologically unrelated plasmids (p<0.001) (**Table 2; Supplementary Table S3**). Between the three comparison methods, the number of SNP differences and the Roary phylogenetic distance showed the highest degree of correlation with a Spearman’s rank correlation coefficient of 0.820 (p < 0.001). The Spearman’s rank correlation coefficients between the Stoesser similarity index and the number of SNP differences or Roary phylogenetic distance were −0.65 (p < 0.001) and −0.77 (p < 0.001), respectively.

**Table 2.** Number of pairwise comparisons, SNP differences, genes variably present or present, and combined number of SNP differences and variable gene presence between the plasmids.

|  | SNP differences |  | Phylogenetic distance using roary |  | Stoesser-similarity index |  |
| --- | --- | --- | --- | --- | --- | --- |
|  | Related | Unrelated | Related | Unrelated | Related | Unrelated |
| n of pairwise comparisons | 15 | 120 | 15 | 120 | 15 | 120 |
| Median | 0 | 29 | 0.00 | 0.37 | 1.00 | 0.90 |
| Range | 0 - 1 | 0 - 674 | 0.00 - 0.06 | 0.00 - 1.69 | 1.00 - 0.99 | 1 - 0.51 |

When setting the threshold at the minimal value (or maximal value for the Stoesser similarity index) that classified 100% of epidemiologically related plasmid pairs as genetically related, the percentage of presumed epidemiologically unrelated plasmid pairs that were classified as epidemiologically related was 12.5% with the SNP differences approach (**Figure 1a**), 6.7% with the Roary phylogenetic distance approach (**Figure 1b**), and 20.8% for the Stoesser-similarity index approach (**Figure 1c**).

**Figure 1.**
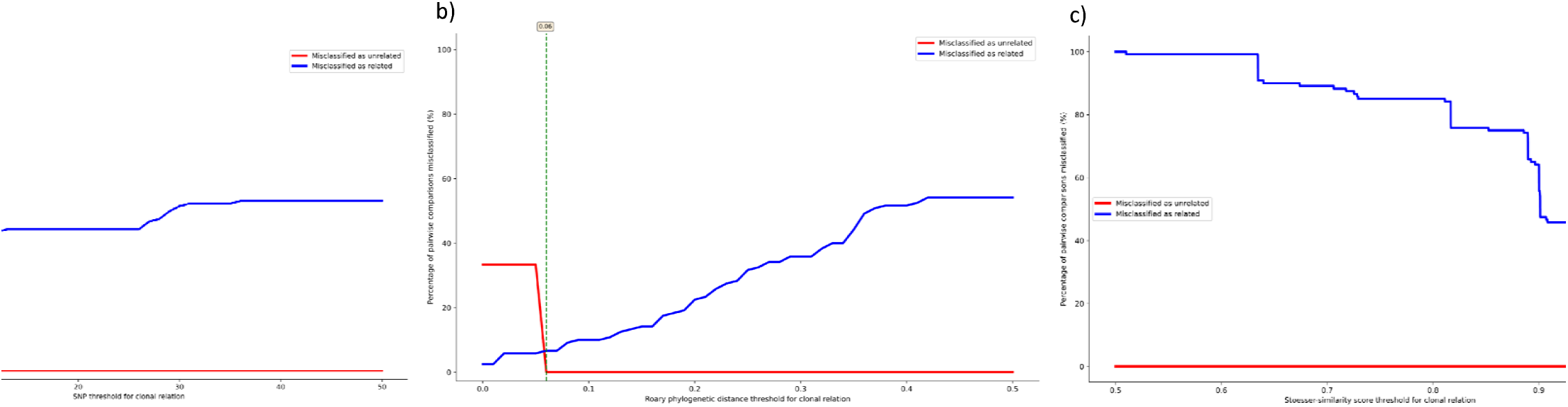
Percentage of pairwise comparisons of epidemiologically related and unrelated plasmids misclassified as either genetically related (blue) or genetically unrelated (red) using different thresholds of **a)** number of SNP differences **b)** Roary phylogenetic distance **c)** Stoesser-similarity index. Green dotted line (number in the text box): minimal threshold value (or maximal value for the Stoesser-similarity index) that classified 100% of epidemiologically related plasmid pairs as genetically related.

## Discussion

Based on our results, epidemiologically related plasmids can be distinguished from unrelated plasmids based on genetic similarity. However, when using a genetic similarity threshold that classified 100% of epidemiologically related plasmid pairs as genetically related, the percentages of plasmids misclassified as epidemiologically related based on the sequence data ranged from 6.7% to 20.8% depending on the comparison method. The phylogenetic distance determined using Roary showed the highest discriminatory power between the epidemiologically related and unrelated plasmids.

Molecular plasmid typing has been used to demonstrate the transmission of *bla*_KPC_-containing plasmids between bacterial isolates and patients (^13,14^). However, previous studies did not include similar plasmids of unrelated patients in the analysis. Our findings on the degree of sequence similarity between *bla*_KPC_-containing plasmids isolated from epidemiologically related and unrelated patients can be used to reveal a possible horizontal transfer of *bla*_KPC_-containing IncFII(k2)-IncFIB(pQiL) plasmids in hospital outbreaks. Despite this, when using the lowest (or highest for the Stoesser-similarity index) threshold for clonal relatedness that classified 100% of epidemiologically related plasmid pairs as genetically related, the percentage of presumed epidemiologically unrelated plasmid pairs that were classified as genetically related was higher in *bla*_KPC-_-containing IncFII(k2)-IncFIB(pQiL) plasmids as compared to the percentages described when typing the bacterial chromosome in a previous study (^22^). This relatively high percentage of misclassifications compared to bacterial chromosome typing was present in all three comparison methods, suggesting a relatively stable plasmid content. Several other studies have also described a high degree of sequence similarity between *bla*_KPC_ containing plasmids isolated from epidemiologically unrelated patients (^28,30^). However, to the best of our knowledge, this is the first study comparing the sequence similarity between epidemiologically related and unrelated *bla*_KPC_-containing IncFII(k2)-IncFIB(pQiL) plasmids. The limited number of differences between plasmids isolated from epidemiologically unrelated patients observed in this study is also seen for other resistance plasmids (^31,32^). A recent study has stated that OXA-48 containing plasmids from outbreak-related patients could not be distinguished from similar plasmids of non-outbreak related patients based on SNP differences (^31^). Interestingly, other studies found *bla*_KPC_-containing plasmids to be highly variable in their nucleotide sequence within the same outbreak and even within patients (^15,20^). This suggests that the plasmid content could also be highly unstable *in vivo* during hospital outbreaks. Therefore, it could well be that transmission of *bla*_KPC_-containing IncF plasmids within hospital outbreaks cannot be dismissed based on sequence dissimilarity between the different plasmids investigated.

This study included bacterial isolates encompassing three different species isolated from an extensively described outbreak with clear epidemiological links between the different outbreak patients. Moreover, the epidemiologically related *bla*_KPC_-containing IncFII(k2)-IncFIB(pQiL) plasmids were compared to a set of similar unrelated plasmids using both a SNP and gene-by-gene based approach in a setting where *bla*_KPC_-containing Enterobacteriaceae are non-endemic. This limits the possibility of falsely classifying *bla*_KPC_-containing plasmids as epidemiologically related. Furthermore, all epidemiologically related *bla*_KPC_-containing IncFII(k2)-IncFIB(pQiL) plasmid sequences in this study were single circular contigs in the assembly graph.

This study has several limitations. Only *bla*_KPC_-containing IncFII(k2)-IncFIB(pQiL) plasmid sequences of six isolates in one outbreak were included. Therefore, it remains unknown whether plasmids belonging to other incompatibility groups and containing other resistance genes also show a high degree of sequence similarity for epidemiologically related plasmids. Moreover, studies that include more epidemiologically related and unrelated *bla*_KPC_-containing IncFII(k2)-IncFIB(pQiL) plasmids are needed to confirm our findings.

To conclude, *bla*_KPC_-containing IncFII(k2)-IncFIB(pQiL) plasmids isolated from epidemiologically related and unrelated Enterobacteriaceae show a high degree of sequence similarity. Judgements on the horizontal transfer of these plasmids during hospital outbreaks based on genetic identity should be made with caution. The phylogenetic distance as determined using Roary showed the highest degree of discriminatory power between the epidemiologically related and unrelated plasmids

## Supplementary material

Supplementary table S1, S2, S3

## Data bibliography

Short-and long-read sequence data included in this study are available from the publicly available European Nucleotide Archive of the European Bioinformatics Institute under study accession number: PRJEB41009. Accession numbers of the plasmid sequences extracted from the GenBank are listed in **Table 1**.

## Abbreviations

KPC: *Klebsiella pneumoniae* carbapenemase
SNP: Single nucleotide polymorphism
wgMLST: Whole-genome MLST
Stoesser-similarity index: similarity index as described by Stoesser et al.

